# *Theobroma grandiflorum* breeding optimization based on repeatability, stability and adaptability information

**DOI:** 10.1101/2021.06.01.446536

**Authors:** Saulo Fabrício da Silva Chaves, Rafael Moysés Alves, Rodrigo Silva Alves, Alexandre Magno Sebbenn, Marcos Deon Vilela de Resende, Luiz Antônio dos Santos Dias

**Affiliations:** Universidade Federal de Viçosa, Viçosa, Minas Gerais, Brazil; Embrapa Amazônia Oriental, Belém, Pará, Brazil; Instituto Nacional de Ciência e Tecnologia do Café, Lavras, Minas Gerais, Brazil; Instituto Florestal de São Paulo, São Paulo, São Paulo, Brazil; Embrapa Café, Viçosa, Minas Gerais, Brazil

**Keywords:** repeated measures, genotype x environment interaction, genotype x measure interaction, REML/BLUP, fruit tree breeding

## Abstract

The cultivation of *Theobroma grandiflorum* in the Brazilian Amazon is mainly conducted by family farmers who use a range of different management strategies. Thus, breeding programs of the species must address the challenge of developing cultivars that are adapted to and stable in a variety of cultivation environments. In this context, this study aimed to estimate the optimum number of harvests for genetic selection of *T. grandiflorum* progenies and identify the most promising ones in terms of productivity, stability, and adaptability. The trials were implemented in three environments, using a randomized complete block design, with 25 full-sib progenies, five replications, and three plants per plot. The traits mean number of fruits/plant, mean fruit production/plant, and rate of infection with witches’ broom (*Moniliophthora perniciosa*) were evaluated over 11 harvests. The Restricted Maximum Likelihood/Best Linear Unbiased Prediction (REML/BLUP) mixed model method was used to estimate genetic parameters and predict genetic values, which were then applied to assess stability and adaptability. The results show that there is genetic variability among the studied *T. grandiflorum* progenies and that accurate genetic selection aiming at recombination is effective after three harvests, for recombination, or eleven harvests for identification of recommended progenies. Six progenies were selected that met the requirements for productivity, stability, and adaptability to different cultivation environments. These results can be used to optimize and advance *T. grandiflorum* breeding programs.

## Introduction

The allogamous tree *Theobroma grandiflorum* (Willd. Ex Spreng.) Schum. (Malvaceae family), commonly known as cupuassu tree, is native to Southeast Pará and Northwest Maranhão States in the Brazilian Amazon [1]. Due to the movement of indigenous peoples throughout the interior of the Amazon region, the species is now dispersed across all Amazonian states [2], and plantations of *T. grandiflorum* have been established in 97 (67%) of the 144 municipalities in Pará State [3]. These plantations are generally small-scale seed orchards of less than one hectare, planted by family farmers [4]. The expansion of the crop and its adaptation to different environments in Pará is an indicator of the genetic plasticity of the species [2].

The economic importance of *T. grandiflorum* has grown in recent years as the main products derived from the tree, including its seeds and the pulp covering them, have attracted increased attention in national and international markets [5]. The pulp, with high acidity and strong aroma, is used to produce juices, sweets, and jellies, among other food products [6]. The almonds, which have antioxidant properties, are used in the pharmaceutical and cosmetic industries [7], as well as to produce cupuassu chocolate, a product known as “cupulate” [8]. The municipality of Tomé Açu, Northeast Pará, was a pioneer in the cultivation of this fruit tree. The region has become a model for production as farmers have organized an agricultural cooperative that processes all cupuassu products, which is essential for expanding its production and use in the region [9]. To ensure the development and sustainability of the crop, communities have continuously sought research support, particularly in terms of developing varieties that are well adapted to local conditions.

At the end of the 1980s, Embrapa Amazonia Oriental initiated a *T. grandiflorum* breeding program and developed genetic resources to produce genotypes with high levels of fruit production and tolerance to the fungus *Moniliophthora perniciosa*, etiological agent of the witches’ broom disease, a pathogen that can affect the cultivation of all species of the *Theobroma* genus, including *T. grandiflorum* and *T. cacao* [10-11]. However, the previously developed genotypes have inconsistent fruit production when subjected to different environments.

Currently, the Restricted Maximum Likelihood/Best Linear Unbiased Prediction (REML/BLUP) mixed model method is the standard for analyses of genotype x environment (GE) interaction [12-13] and repeated measures [14-15]. There are numerous reasons for its use, including the fact that it enables the simultaneous estimates of variance components and prediction of genetic values. The method also deals well with unbalanced data, describes the heterogeneity of genetic covariances and residual variances across environments, and models spatial trends [16].

The evaluation of different genotypes in a variety of environments enables the quantification of the GE interaction effect [17] and the analysis of genotypic stability and adaptability [18]. Understanding stability and adaptability enables the identification of productive, stable, and adaptable genotypes [19]. However, evaluating GE interaction is one of the most costly aspects of a breeding program [20], especially for perennial fruit trees such as *T. grandiflorum*, where the breeding cycle can last up to 15 years [21]. This may explain why studies on GE interaction in *T. grandiflorum* are extremely rare.

Variation throughout years can create different environments, which, in turn, will influence genotypes differently [22]. The evaluation of genotypes across several harvests is crucial in perennial fruit trees as it enables the quantification of the genotype x measurement (GM) interaction effect and estimates of the repeatability coefficient to determine the optimal number of harvests necessary to conduct effective genetic selection [15, 17].

In this context, this study aimed to estimate the optimum number of harvests for genetic selection of *T. grandiflorum* progenies and identify the most promising progenies in terms of productivity, stability, and adaptability.

## Material and methods

### Experimental data

Full-sib *T. grandiflorum* progeny tests were established in three farms in Northeastern Pará State, Brazil; two located in the municipality of Tomé Açu and one in the municipality of São Francisco do Pará, approximately 210 km apart. The three environments represent a sample of the different cultivation systems used to produce *T. grandiflorum* in Pará. This experimental system enables the evaluation and selection of genotypes for conditions similar to those in which they are often cultivated. The differences between the three environments are mainly the different cropping systems used for each trial, in terms of temporary and definitive shading or full sun, and spacing.

Each *T. grandiflorum* progeny test was installed in consortium with other tree species, all of which were planted in February 2005. The field arrangement affected conditions of luminosity and competition over and under the soil. In trial 1, *T. grandiflorum* was maintained in shade during the productive phase, while in the other two trials (trials 2 and 3) the trees were kept in full sun (Table 1). In trial 1, *T. grandiflorum* progenies were part of an agroforestry system (AFS), together with *Passiflora edulis* Sims. (passion fruit) and *Swietenia macrophylla* King. (Brazilian mahogany), at initial densities of 400, 800, and 100 plants/ha, respectively. After the third year, the passion fruit was removed from the AFS as it had completed its cycle. Therefore, through all production stages, *T. grandiflorum* was shaded with *S. macrophylla*. Trial 2 was also installed as an AFS and consisted of *T. grandiflorum* progenies, *Piper nigrum* L. (Black pepper), and *Bertholletia excelsa* Bonpl. (Brazil nut), at densities of 303, 1800, and 75 plants/ha, respectively. As *B. excelsa* developed a very prolific crown, producing too much shade for *T. grandiflorum* in the first years, it was removed from the AFS in the fifth year. *Piper nigrum* cultivation occurred over the first seven years of the trial. Thus, after *P. nigrum* tree mortality in the seventh year, *T. grandiflorum* was left in full sun. In trial 3, *T. grandiflorum* progenies were intercropped with *Musa* spp. (banana tree), both with a density of 400 plants/ha. As in trial 2, after the fifth year, the *Musa* spp. was removed from the AFS, with *T. grandiflorum* progenies remaining in full sun until the end of the study (Table 1). It is important to highlight that these different field arrangements, involving full sun and temporary and definitive shading, were designed to reproduce a similar environment to what farmers cultivate cupuassu tree in the state of Pará.

**Table 1.**
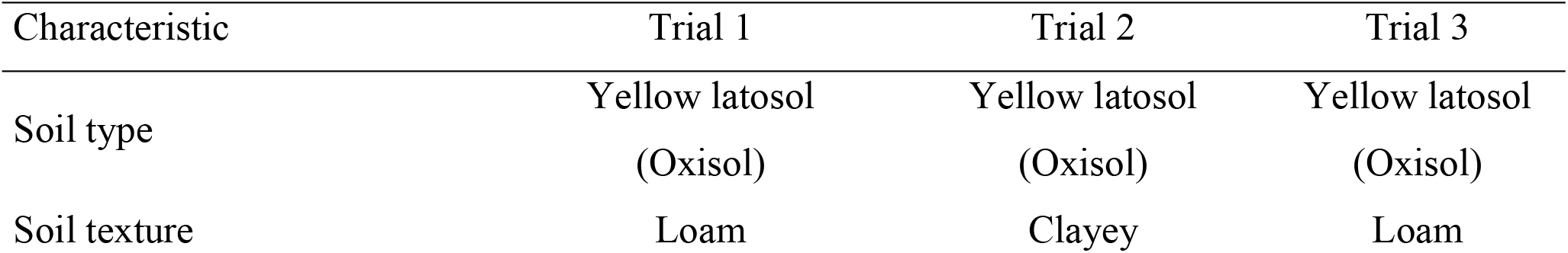
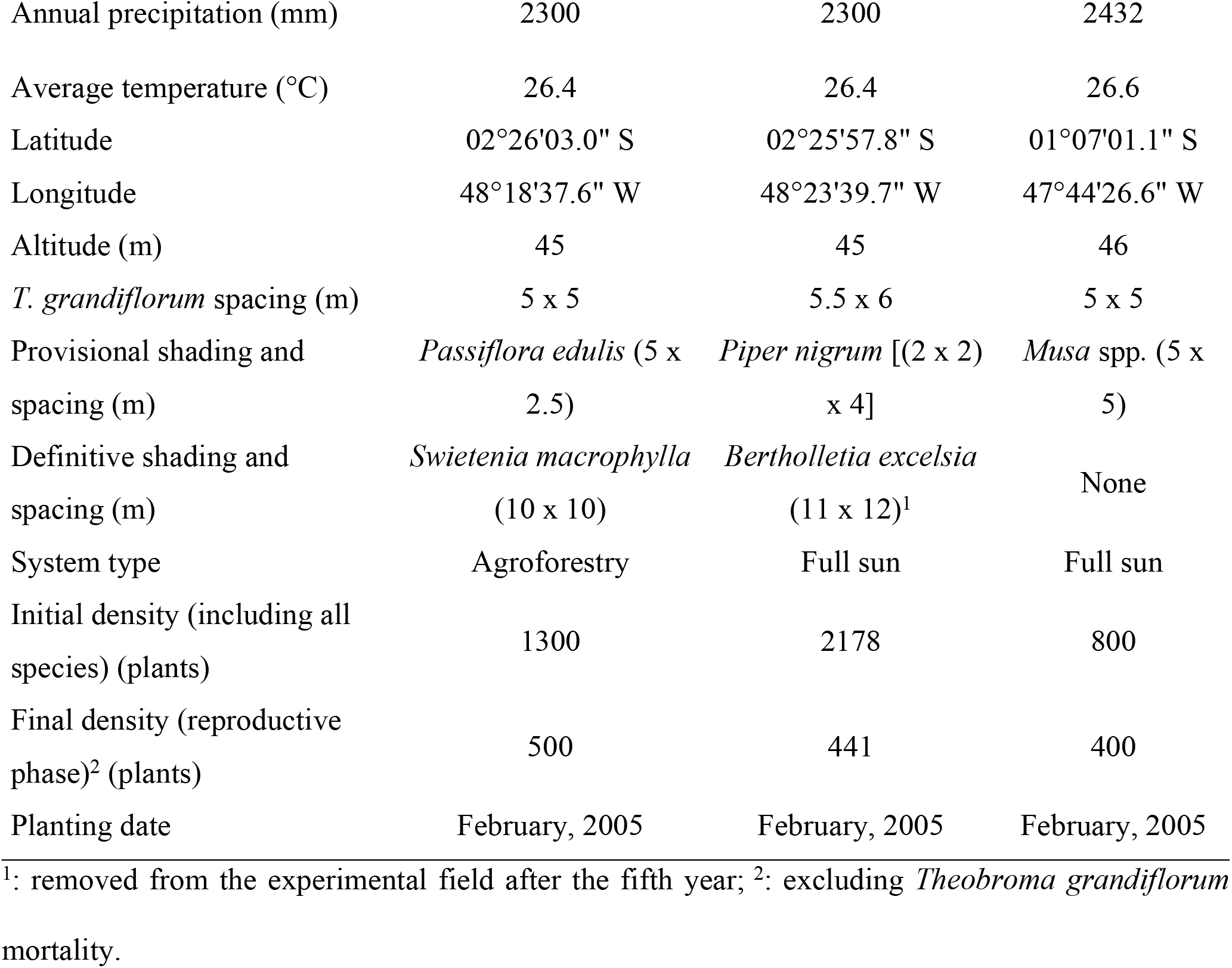
Characteristics of the trials with 25 full-sib *Theobroma grandiflorum* progenies in Northeast Pará State, Brazil.

| Characteristic | Trial 1 | Trial 2 | Trial 3 |
| --- | --- | --- | --- |
| Soil type | Yellow latosol<br>(Oxisol) | Yellow latosol<br>(Oxisol) | Yellow latosol<br>(Oxisol) |
| Soil texture | Loam | Clayey | Loam |
| Annual precipitation (mm) | 2300 | 2300 | 2432 |
| Average temperature (°C) | 26.4 | 26.4 | 26.6 |
| Latitude | 02°26'03.0" S | 02°25'57.8" S | 01°07'01.1" S |
| Longitude | 48°18'37.6" W | 48°23'39.7" W | 47°44'26.6" W |
| Altitude (m) | 45 | 45 | 46 |
| *T. grandiflorum* spacing (m) | 5 x 5 | 5.5 x 6 | 5 x 5 |
| Provisional shading and spacing (m) | *Passiflora edulis* (5 x 2.5) | *Piper nigrum* [(2 x 2) x 4] | *Musa* spp. (5 x 5) |
| Definitive shading and spacing (m) | *Swietenia macrophylla* (10 x 10) | *Bertholletia excelsia* (11 x 12)<sup>1</sup> | None |
| System type | Agroforestry | Full sun | Full sun |
| Initial density (including all species) (plants) | 1300 | 2178 | 800 |
| Final density (reproductive phase)<sup>2</sup> (plants) | 500 | 441 | 400 |
| Planting date | February, 2005 | February, 2005 | February, 2005 |

<sup>1</sup>: removed from the experimental field after the fifth year; <sup>2</sup>: excluding *Theobroma grandiflorum* mortality.

The progeny test, carried out in three trials, was established using a randomized complete block design, with 25 *T. grandiflorum* full-sib progenies, five replications, and three plants per plot. The 25 full-sib progenies were obtained through controlled pollination. Phenotypic data were measured over 11 consecutive annual harvests, based on a total plot. The harvest opening coincides with the beginning of the rainy season and extends over the entire period of about six months. Therefore, each harvest was divided into four evaluations with 45-day intervals between them. Response variables included the mean number of fruits/plant (NF) and mean fruit production (kg/plant), obtained by multiplying the NF by the average weight of the fruit of each genotype. We also assessed tolerance to witches’ broom disease (*M. perniciosa*) based on the rate of symptomatic plants per progeny (WB, %). Plants were deemed symptomatic when at least one branch presented misshapen phyllotaxis, compared to a normal branch, and after one month appeared desiccated [23]. For selection, a tolerance threshold of 30% was adopted as the maximum rate of symptomatic plants per progeny.

### Statistical analyses

Due to the unbalance caused by the mortality of some trees, which is common in long-term trials involving perennial crops, the mixed model method was adopted for statistical analysis. With such an approach, the variance components and genetic parameters are estimated by the restricted maximum likelihood (REML) [24], and genetic values are predicted by the best linear unbiased prediction (BLUP) [25]. The mixed linear model associated with the analysis of progeny, with a complete randomized block design, three locations, at the plot level, and with repeated measures, is defined as:

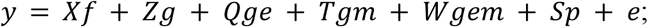

where, *y* is the vector of phenotypic data; *f* is the vector of the effects of the combination repetition-environment-measurement (assumed to be fixed), added to the general mean; *g* is the vector of genotypic effects (assumed to be random), *g* ∼ NID 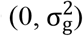, where 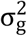 is the genotypic variance; *ge* is the vector of genotypes × environments (GE) interaction effects (assumed to be random) *ge* ∼ NID 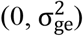, where 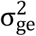 is the GE interaction variance; *gm* is the vector of genotypes × measurements (GM) interaction effects (random) *gm* ∼ NID 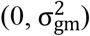, where 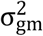 is the GM interaction variance; *gem* is the vector of the triple genotypes x environments x measurements (GEM) interaction effects (random) *gem* ∼ NID 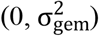, where 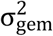 is the GEM interaction variance; *p* is the vector of the permanent plot effects within locations (assumed to be random) *p* ∼ NID 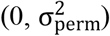, where 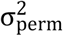 is the permanent plot effect variance; and *e* is the vector of residuals (random) *e* ∼ NID 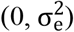, where 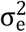 is the residual variance. The capital letters (*X, Z, Q, T, W* e *S*) represent the incidence matrices for the correspondents’ effects.

The significance of the random effects of the statistical model was tested by the likelihood ratio test (LRT), given by the following equation [26]:

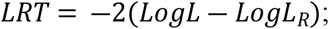

where, *LogL* is the logarithm of the maximum point of the residual likelihood function (*L*) of the complete model; and *LogL*_*R*_ is the logarithm of the maximum point of the residual likelihood function (*L*_*R*_) of the reduced model (without the effect under test). The chi-square statistic with one degree of freedom and a probability level equal to 1% was used to test the LRT significance.

From the variance components (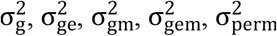 and 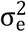), we estimated [14]:

Phenotypic variance: 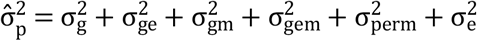;

Mean phenotypic variance: 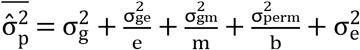,

where e, m, and b are the number of environments, measurements, and blocks, respectively;

Individual broad-sense heritability: 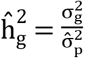;

Mean broad-sense heritability: 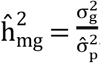;

Accuracy of genotype selection: 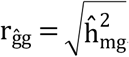;

Coefficient of determination of GE interaction effects: 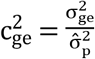;

Coefficient of determination of GM interaction effects: 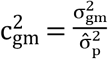;

Coefficient of determination of GEM interaction effects: 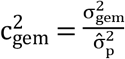;

Coefficient of determination of plot effects: 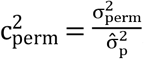;

Coefficient of individual repeatability: 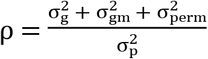;

Genotypic correlation among environments: 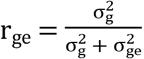;

Genotypic correlation among measurements: 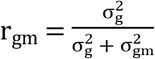; and

Genotypic correlation among environments and measurements: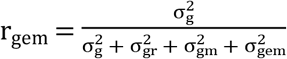.

To determine the optimal number of harvests for genetic selection, accuracy was calculated considering the use of *m* harvests (r_m_) [14]:

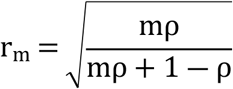

We also considered the efficiency (E) of the use of *m* harvests in relation to the use of only one harvest [14]:

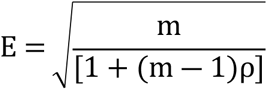

To select progenies with greater adaptability, stability, and productivity, the Harmonic Mean of Relative Performance of Genotypic Values (HMRPGV) method was used. This method provides a genotypic value that is affected negatively by instability and positively by adaptability [27]:

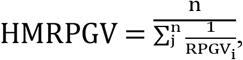

where, *n* is the number of environments; *RPGV*_*ij*_ is the Relative Performance of Genotypic Values, estimated as

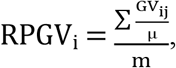

where, *GV*_*ij*_ is the genotypic value of the *i*^*th*^ genotype in the *j*^*th*^ environment, and *μ* is the phenotypic mean of the *j*^*th*^ environment. All statistical analyses were performed using the SELEGEN REML/BLUP software [16].

## Results

The genotypic effects were significant for both the mean number of fruits/plant and mean fruit production/plant, indicating genetic variability among progenies. Furthermore, the presence of GE and GEM interaction was verified for these traits. As expected for polygenic traits, there was a high level of influence of uncontrolled environmental factors, which is reflected in the residual variance that corresponds to the largest proportion of phenotypic variance (Table 2).

**Table 2.**
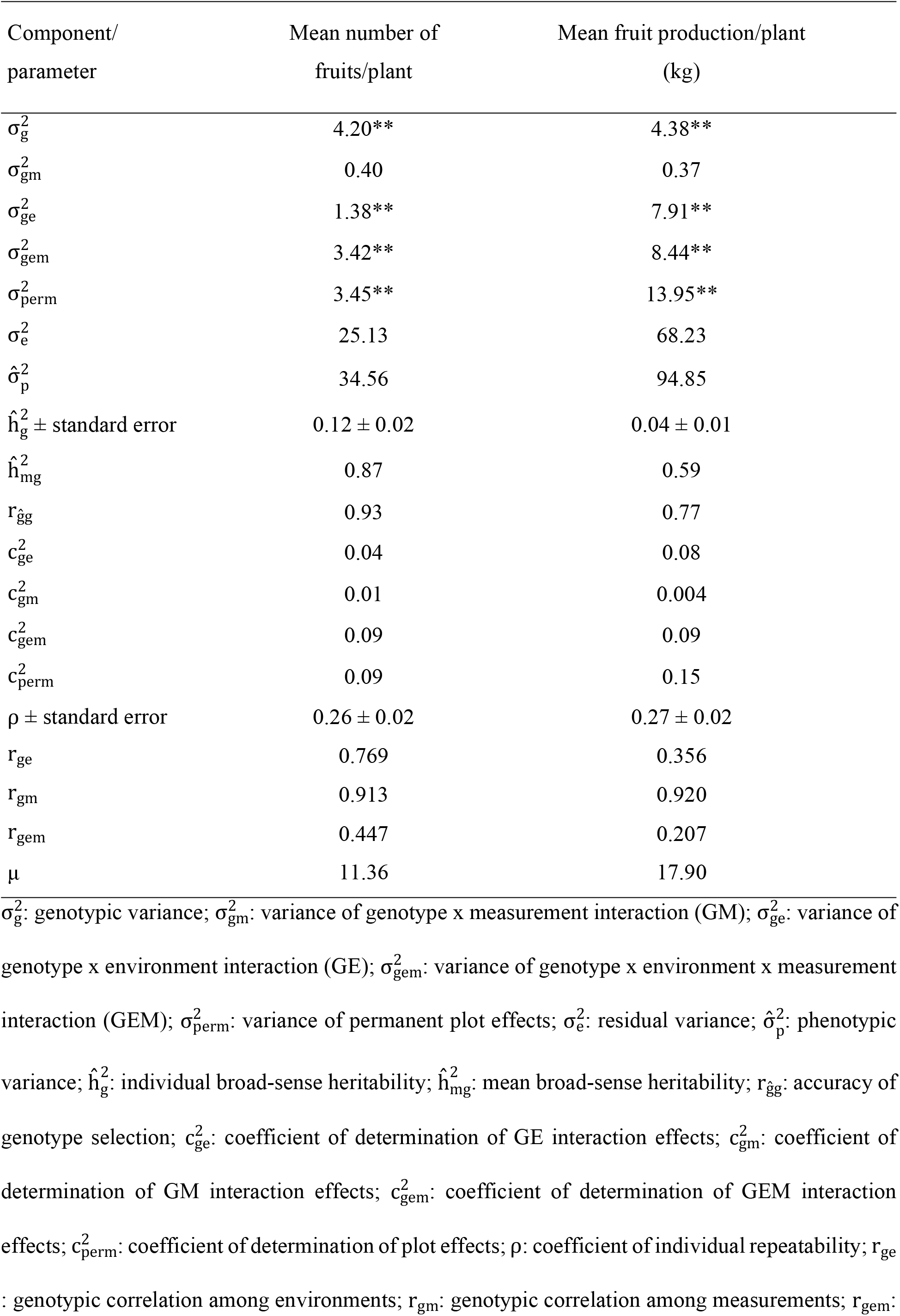
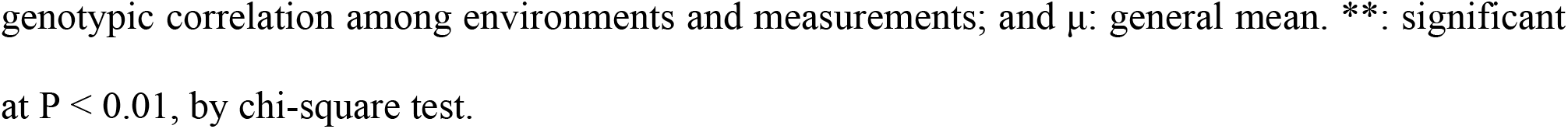
Estimates of the components of variance and genetic parameters for mean number of fruits/plant and mean fruit production/plant, evaluated in 25 full-sib progenies of *Theobroma grandiflorum*.

| Component/<br>parameter | Mean number of<br>fruits/plant | Mean fruit production/plant<br>(kg) |
| --- | --- | --- |
| $\sigma_g^2$ | 4.20** | 4.38** |
| $\sigma_{gm}^2$ | 0.40 | 0.37 |
| $\sigma_{ge}^2$ | 1.38** | 7.91** |
| $\sigma_{gem}^2$ | 3.42** | 8.44** |
| $\sigma_{perm}^2$ | 3.45** | 13.95** |
| $\sigma_e^2$ | 25.13 | 68.23 |
| $\hat{\sigma}_p^2$ | 34.56 | 94.85 |
| $\hat{h}_g^2 \pm$ standard error | $0.12 \pm 0.02$ | $0.04 \pm 0.01$ |
| $\hat{h}_{mg}^2$ | 0.87 | 0.59 |
| $r_{\hat{g}g}$ | 0.93 | 0.77 |
| $c_{ge}^2$ | 0.04 | 0.08 |
| $c_{gm}^2$ | 0.01 | 0.004 |
| $c_{gem}^2$ | 0.09 | 0.09 |
| $c_{perm}^2$ | 0.09 | 0.15 |
| $\rho \pm$ standard error | $0.26 \pm 0.02$ | $0.27 \pm 0.02$ |
| $r_{ge}$ | 0.769 | 0.356 |
| $r_{gm}$ | 0.913 | 0.920 |
| $r_{gem}$ | 0.447 | 0.207 |
| $\mu$ | 11.36 | 17.90 |

$\sigma_g^2$ : genotypic variance; $\sigma_{gm}^2$ : variance of genotype x measurement interaction (GM); $\sigma_{ge}^2$ : variance of genotype x environment interaction (GE); $\sigma_{gem}^2$ : variance of genotype x environment x measurement interaction (GEM); $\sigma_{perm}^2$ : variance of permanent plot effects; $\sigma_e^2$ : residual variance; $\hat{\sigma}_p^2$ : phenotypic variance; $\hat{h}_g^2$ : individual broad-sense heritability; $\hat{h}_{mg}^2$ : mean broad-sense heritability; $r_{\hat{g}g}$ : accuracy of genotype selection; $c_{ge}^2$ : coefficient of determination of GE interaction effects; $c_{gm}^2$ : coefficient of determination of GM interaction effects; $c_{gem}^2$ : coefficient of determination of GEM interaction effects; $c_{perm}^2$ : coefficient of determination of plot effects; $\rho$ : coefficient of individual repeatability; $r_{ge}$ : genotypic correlation among environments; $r_{gm}$ : genotypic correlation among measurements; $r_{gem}$ :
genotypic correlation among environments and measurements; and $\mu$ : general mean. \*\*: significant at $P < 0.01$ , by chi-square test.

Only mean broad-sense heritability of genotypes 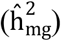 can be considered high. Both mean and individual broad-sense heritability 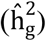 were higher for mean number of fruits/plant than for mean fruit production/plant. Selective accuracy (r_ĝg_) followed the same pattern. The coefficient of repeatability (ρ) showed similar magnitudes for mean number of fruits/plant and mean fruit production/plant, a positive aspect in selection when considering both traits simultaneously since the optimal number of measurements will coincide. With the use of a greater number of measurements, compared to only one measurement, the selective accuracy increases (Fig 1). With the use of data from three harvests, the selective accuracy exceeds 0.70; with the use of 11 harvests, the selective accuracy exceeds 0.90 (Fig 1A). The efficiency associated with the use of *m* measures indicates smaller increments as the number of harvests increases (Fig 1B). These increments become almost constant as of the ninth harvest, with only 2% increase compared to the previous harvest.

**Fig 1.**
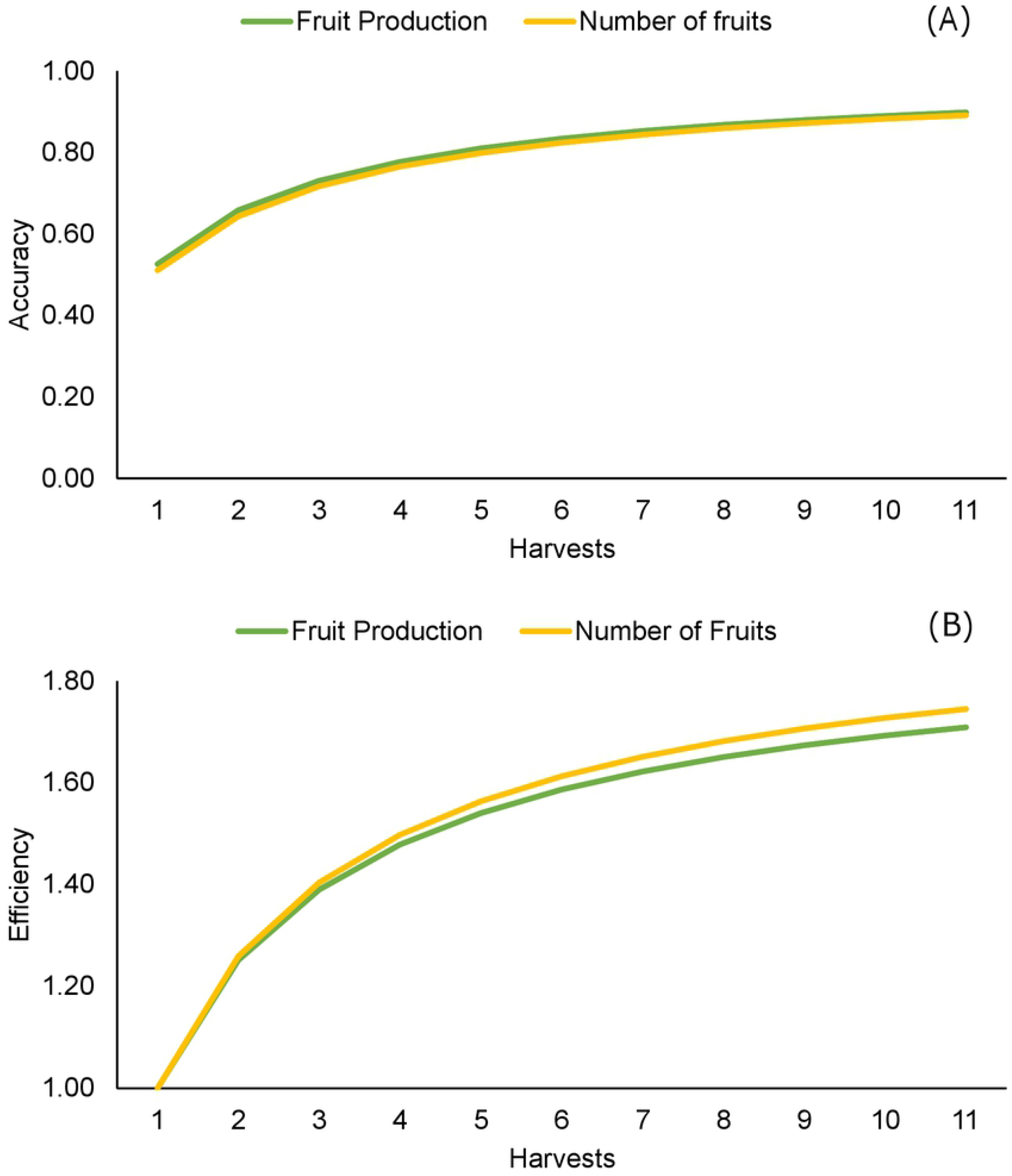
Accuracy (A) and efficiency (B) of selection as a function of the number of measures.

The coefficient of determination of GE interaction effects 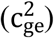, GM interaction effects 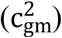, and plot effects 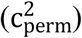 were all low for both traits (ranging from 0.004 to 0.15). The genotypic correlations across environments (r_ge_) and environments and measurements (r_gm_) were higher for mean number of fruits/plant, while the genotypic correlation across measurements (r_gm_) was slightly higher for mean fruit production/plant.

For mean number of fruits/plant, the coincidence was high (90%) between HMRPGV, genotypic values (μ + g), and genotypic values plus the mean effect of the GE interaction (μ + g + gem), considering the selection of the 10 best progenies. This demonstrates that the most productive progenies are also the most adapted and stable. The selection of the ten best progenies (36, 37, 11, 5, 49, 23, 6, 43, 19, and 25) provided a predicted selection gain of 2.61 fruits/plant (∼ 23% compared to the general mean) (Table 3).

**Table 3.** Genotypic values (μ + g), genotypic values plus the average GE interaction effect (μ + g + gem), Harmonic Mean of the Relative Performance of Genotypic Values multiplied by the General Mean (HMRPGV *μ), and genetic gain with selection (Gain), for mean number of fruits/plant, mean fruit production/plant, and incidence rate of witches’ broom (WB, %), evaluated in 25 full-sib progenies (Prog) of *Theobroma grandiflorum*.

| Mean number of fruits/plant |  |  |  |  | Mean fruit production/plant (kg) |  |  |  |  |  |
| --- | --- | --- | --- | --- | --- | --- | --- | --- | --- | --- |
| Prog<sup>1</sup> | ($\mu + g$) | ($\mu + g + \text{gem}$) | (HMRPGV * $\mu$) | Gain | Prog<sup>1</sup> | ($\mu + g$) | ($\mu + g + \text{gem}$) | (HMRPGV * $\mu$) | Gain | WB (%) |
| 36 | 15.25 | 15.67 | 15.70 | 3.89 | 36 | 21.13 | 23.08 | 23.10 | 3.23 | 37.3 |
| **37** | 14.61 | 14.96 | 15.03 | 3.57 | **5** | 19.98 | 21.24 | 21.58 | 2.66 | 11 |
| **11** | 13.71 | 13.96 | 13.85 | 3.16 | **37** | 19.84 | 21.01 | 20.97 | 2.42 | 25.8 |
| **5** | 13.49 | 13.73 | 13.63 | 2.91 | **11** | 19.41 | 20.32 | 20.57 | 2.09 | 11 |
| **49** | 13.09 | 13.28 | 13.20 | 2.67 | **23** | 19.56 | 20.55 | 20.43 | 2.23 | 29 |
| **23** | 12.95 | 13.13 | 13.11 | 2.49 | **49** | 19.14 | 19.89 | 20.10 | 1.95 | 20 |
| 6 | 12.25 | 12.35 | 12.31 | 2.26 | **21** | 18.54 | 18.93 | 18.92 | 1.76 | 8.3 |
| 43 | 12.13 | 12.21 | 11.97 | 2.08 | 6 | 18.41 | 18.72 | 18.81 | 1.61 | 32.7 |
| 19 | 11.54 | 11.56 | 11.59 | 1.46 | **25** | 18.01 | 18.08 | 17.91 | 1.24 | 23.5 |
| **25** | 11.57 | 11.59 | 11.57 | 1.57 | 38 | 17.84 | 17.81 | 17.83 | 1.14 | 33.8 |
| 38 | 11.58 | 11.60 | 11.56 | 1.71 | 52 | 18.35 | 18.63 | 17.75 | 1.48 | 37.5 |
| 52 | 11.62 | 11.65 | 11.55 | 1.88 | 43 | 18.18 | 18.35 | 17.66 | 1.36 | 69.8 |
| 8 | 11.19 | 11.17 | 11.18 | 1.33 | 9 | 17.71 | 17.60 | 17.46 | 1.03 | 37.8 |
| 4 | 11.10 | 11.07 | 11.06 | 1.22 | 8 | 17.66 | 17.51 | 17.38 | 0.94 | 8.7 |
| 1 | 10.92 | 10.87 | 10.80 | 1.11 | 22 | 17.42 | 17.13 | 17.01 | 0.70 | 37.7 |
| **21** | 10.75 | 10.69 | 10.72 | 0.91 | 20 | 17.55 | 17.34 | 16.97 | 0.86 | 19.7 |
| 9 | 10.80 | 10.74 | 10.67 | 1.00 | 4 | 17.20 | 16.79 | 16.69 | 0.63 | 22.3 |
| 20 | 10.37 | 10.26 | 10.29 | 0.80 | 19 | 17.11 | 16.64 | 16.62 | 0.55 | 15.8 |
| 22 | 10.15 | 10.02 | 10.06 | 0.70 | 13 | 17.49 | 17.25 | 16.43 | 0.78 | 72.8 |
| 17 | 9.90 | 9.74 | 9.78 | 0.49 | 1 | 17.11 | 16.64 | 16.13 | 0.49 | 28.7 |
| 13 | 9.96 | 9.80 | 9.54 | 0.59 | 17 | 16.56 | 15.76 | 15.66 | 0.40 | 37.5 |
| 30 | 9.51 | 9.31 | 9.25 | 0.39 | 30 | 16.45 | 15.57 | 15.59 | 0.32 | 44.3 |
| 40 | 8.99 | 8.73 | 8.76 | 0.27 | 40 | 15.97 | 14.82 | 14.44 | 0.22 | 26.7 |
| 29 | 8.69 | 8.40 | 8.42 | 0.15 | 29 | 15.70 | 14.37 | 14.41 | 0.12 | 17 |
| 28 | 7.85 | 7.47 | 7.42 | 0.00 | 28 | 15.09 | 13.40 | 13.23 | 0.00 | 13.5 |
| Means | 11.36 | 11.36 | 11.32 | 1.54 |  | 17.90 | 17.90 | 17.75 | 1.21 | 28.89 |

'progenies are listed in decreasing order in terms of results for mean number of fruits/plant and mean fruit production/plant, respectively. Selected progenies are in bold.

For mean fruit production/plant (kg), which is the trait of greatest economic importance, there was an 80% coincidence between μ + g, μ + g + gem, and HMRPGV. Progenies 56 and 43, despite having higher values for μ + g and μ + g + gem compared to genotypes 25 and 38 (ranked 9^th^ and 10^th^, respectively), did not show satisfactory stability. The selection of the ten best progenies (36, 5, 37, 11, 23, 49, 21, 6, 25, and 38) led to a predicted selection gain of 2.03 kg per plant (∼ 11.3% compared to the general mean; Table 3). However, for genetic selection, the incidence rate of witches’ broom must be taken into account. Considering a rate of 30% symptomatic plants per progeny as a tolerance threshold, progenies 36, 6, and 38 should not be selected. Thus, the choice of the seven remaining progenies (5, 37, 11, 23, 49, 21, and 25, in bold) provided a predicted selection gain of 2.05 kg (∼ 11.4% in relation to the general mean).

## Discussion

LRT shows genetic variability among progenies for both the traits mean number of fruits/plant and mean fruit production/plant. Although the GE and GEM interactions were significant, the genotypic correlation among environments (r_ge_) was high (0.77) only for mean number of fruits/plant, according to the classification proposed by [28]. These results indicate that for this trait the performance of progenies is moderately similar among the trials and some of the same progenies can be selected for them all.

The significance of the GE interaction associated with low genotypic correlation among environments (r_ge_= 0.356) for mean fruit production/plant indicate that the cultivation system of *T. grandiflorum* can have a significant influence on the productive performance of different genotypes in different environments. Given that the varied needs of the stakeholders and intended end users must be considered when developing cultivars, it is important to highlight that the vast majority of producers in Northeast Pará use cultivation systems similar to those studied herein. Therefore, it is essential to select genotypes that have satisfactory adaptability, stability, and productivity in a range of management scenarios [29]. In studying *T. cacao*, [30] found that the level of shade, one of the distinguishing characteristics of each environment studied herein, can have an effect on photosynthesis and, thus, productive capacity.

Based on the coefficient of determination of the GE interaction 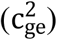 and genotypic correlation among environments (r_ge_), mean fruit production/plant was more heavily influenced by the environment than mean number of fruits/plant. In a previous study [28], the authors highlight that a useful indicator is the ratio between the variances attributed to the GE interaction and the genotype 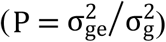. Variables with P < 0.5, as is the case with mean number of fruits/plant (0.33), will not be problematic for selection; while traits with P > 0.5, such as mean fruit production/plant (1.80), tend to generate problems due to the complex GE interaction, especially when the intention is to carry out indirect selection.

Such a pattern is expected since mean fruit production/plant is a quantitative trait composed of two other traits that are also polygenic: the mean number of fruits/plant and the mean fruit weight. Traits of this nature are influenced by the activity of numerous genes, combined with significant environmental effects [31]. This highlights the impact the type of management can have on the phenotypic manifestation of the evaluated traits, especially mean fruit production/plant. This fact, combined with the variability resulting from the species’ self-incompatibility [32], is reflected in uncertainties about the cultivation of genetic materials that have not been evaluated in a range of environments and emphasizes the importance of studies of this nature.

On the other hand, the GM interaction was not significant. According to [15], this result is an indication that there is consistency in the performance of genotypes across the years of evaluation. This was confirmed by the high values of the genotypic correlation through measurements for both mean number of fruits/plant and mean fruit production/plant (> 0.90). The GM interaction is mainly related to the reaction of progenies to climate change and its consequences. The development of improved genetic materials must take into account the variation of the climate between and within years, aiming to increase the resilience of the cultivars offered to producers [33]. The genotypes evaluated in this study fulfil this requirement. When evaluating both components mentioned above jointly through the triple interaction (GEM), differential behavior was observed across trials and years. However, given the non-significance of GM, it appears that most of the GEM interaction is due to the GE interaction.

According to the classification presented by [17], individual broad-sense heritability showed a low magnitude 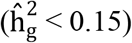 for both traits. In contrast, estimates of mean heritability showed a high magnitude 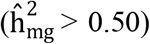. The low individual broad-sense heritability 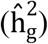 refers to the quantitative nature of both traits, as discussed above, making the process of selection more complex [34]. Through the interpretation of these heritability values, along with the high mean heritability 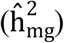 for both traits, we can infer that priority should be given to the selection of progenies, rather than the selection of ortets. This is due to the relationship between heritability and genetic gains with selection, in which the latter is a direct function of the former. Thus, heritability estimates can reveal the selection efficacy, before it is done [35]. Therefore, smaller-scale heritabilities, as observed at the individual level, can jeopardize the genetic gains. A previous study [36] considered this fact to recommend the selection of full-sib families in guava (*Psidium guajava* L.) to achieve greater genetic gains. Another attribute of heritability is related to selection accuracy. According to Resende and Duarte (2007), values above 0.7 are considered high, as is the case for mean fruit production/plant, and above 0.9 they are considered very high, as is the case with mean number of fruits/plant. The selective accuracy, or the correlation between the true and predicted genotypic value, enables us to infer the selection precision [38-39].

Heritability is also related to the repeatability coefficient, which is the maximum value that individual broad-sense heritability can achieve [38]. Estimating the repeatability coefficient using the mixed model method has greater flexibility when compared to ANOVA, as it can be used even when the assumptions required for the analysis of variance are not met [15]. Previous research has highlighted the importance of this parameter for perennial species, which have a long breeding cycle [40]. In that study, the authors obtained a repeatability coefficient of 0.35 for fruit production in *Annona muricata* L., classifying it as moderate, based on [38]. Taking into account this same classification, the estimates of repeatability coefficients for both traits studied herein were of low magnitude (< 0.3), which indicates that the evaluation of several harvests is necessary for genetic selection. To achieve an accuracy of 0.70, the minimum value for selection aiming at recombination [37], data from three consecutive initial harvests are enough, which is consistent with what was found for yield components in *T. cacao* [41, 42]. If the objective is only to recombine and advance the cycle, subsequent measurements are unnecessary as they would offer limited increases in efficiency (Fig 1B) but require more time and incur higher costs related to the measurement of each harvest. If the intention is to identify genetic materials for cultivation, the evaluation of 11 harvests is recommended, since conducting such measurements can offer a selective accuracy of 0.90, a value suggested by [35]. Within the *Theobroma* genus, analyses similar to the present study have only been conducted for *T. cacao* [41, 43] and have offered substantial and fundamental results for breeding programs. For *T. grandiflorum*, the results presented herein will enable optimization of the breeding period, increasing gains by decreasing the time of field assessment from 15 to six years, considering the three years needed for initial establishment (juvenility period). Studies of this nature are rare for *T. grandiflorum*, thus demonstrating the pioneering nature of this work.

The use of HMRPGV proved to be a viable alternative for *T. grandiflorum*, as it allows us to infer the adaptability and stability of genotypic values. We found high levels of coincidence between the best genotypes in HMRPGV and μ + g, what indicates the efficiency of the method [13]. Comparing both studied traits, it is clear that there is no perfect match as to the best genotypes. This is due to the low levels of correlation between the number of fruits and the average weight of fruits, that is, plants that produce heavy fruits and in large quantities are exceptions. The identification of these genotypes is essential for advancing the improvement of the species [21]. In this context, progenies 36, 37, 11, 5, 49, 23, 6, and 25 are the most suitable, as they stand out for both traits simultaneously.

Combining the analysis of productive traits with the resistance to witches’ broom (*M. perniciosa*) disease, progenies 6, 36, and 38 were excluded, as they did not present satisfactory tolerance, and thus, can increase the pathogen pressure on resistant individuals. Thus, of the 25 studied progenies, 5, 37, 11, 23, 49, 21, and 25 were selected. Breeding programs for *T. grandiflorum* must always take into account the plants’ reaction to the fungus since it is the main pathogen that affects the cultivation of both *T. grandiflorum* and *T. cacao* [44]. Therefore, through breeding of the species, genetic materials can be developed that combine high levels of productivity and resistance to *M. perniciosa*, thus reducing production costs related to phytosanitary pruning and the application of fungicide. This, in turn, can mitigate the risks and effects of chemical contamination for humans, animals, and the environment [45].

## Conclusions

Data from three or 11 harvests should be used in selection aiming at recombination or identification of genotypes for selection, respectively.

Seven progenies (5, 37, 11, 23, 49, 21 and 25) were identified and selected with high adaptability, stability, productivity, and resistance to witches’ broom disease.

## Acknowledgements

The authors would like to thank the rural producers Emerson Tokumaru, Paulino Taguchi, Michinori Konagano, Seia Takaki, Elton Takaki, and Maria do Socorro Lima for generously providing their properties to install and conduct the trials. We also thank the agricultural technician José Raimundo Quadros Fernandes for assistance with fieldwork.

## Data Availability Statement

The data that support the findings of this study are included in the supporting information files.

## Author Contributions

All authors have made a significant, direct and intellectual contribution to this paper, and approved it for publication.

## Notes

### Competing Interest Statement

The authors have declared no competing interest.

